# Biological insights from self-perceived facial aging data of the UKBB participants

**DOI:** 10.1101/758854

**Authors:** Simona Vigodner, Raya Khanin

## Abstract

Genetic underpinnings of facial aging are still largely unknown. In this study, we leverage the statistical power of large-scale data from the UK Biobank and perform insilico analysis of genome-wide self-perceived facial aging. Functional analysis reveals significant over-representation of skin pigmentation and immune related pathways that are correlated with facial aging. For males, hair loss is one of the top categories that is highly significantly over-represented in the genetics data associated with self-reported facial aging. Our analysis confirms that genes coding for the extracellular matrix play important roles in aging. Overall, our results provide evidence that while somewhat biased, large-scale self-reported data on aging can be utilized for extracting useful insights into underlying biology, provide candidate skin aging biomarkers, and advance anti-aging skincare.

## Introduction

An increasing number of studies use big data to investigate various aspects of biological and chronological facial aging. These studies used to rely on assessments by specialists such as dermatologists or volunteers. Now, facial features related to aging, as well as perceived age, are more often quantified automatically from photographic or 3D images [1]. Perceived age is defined as the age that a person is visually estimated to be on the basis of physical appearance [2]. Factors that affect perceived ageing include increased sun-damage [3], male pattern baldness [4], gray hair, under eye wrinkles and bags, pigmented spots [5], skin topography (skin micro-texture and wrinkles) [6], decreased facial contrast and reduced skin color uniformity [7]–[8].

One study used photographic images of 264 white Dutch and British twins to investigate the link between perceived age and biological age [9]. The study determined that skin wrinkling, appearance of facial sun-damage, hair graying, and lip height were significantly and independently associated with how old the women looked for their age. The structure of subcutaneous tissue was found to be partly responsible for whether women were perceived to look young. Heritability analyses of the appearance features revealed that perceived age, pigmented age spots, skin wrinkles and the appearance of sun-damage were influenced more or less equally by genetic and environmental factors. Another analysis of photographic assessments, using the VISIA(®) Complexion Analysis System, of 67 pairs of Japanese twins found that skin aging, especially in terms of facial texture, was significantly influenced by environmental factors [10]. A large random sample from the Leiden Longevity Study found that a higher non-fasted IGF-1 level was associated with a lower skin wrinkling grade and tended to associate with a lower perceived age, while higher fasted cortisol and higher glucose levels tended to be associated with a higher perceived age [11], [12]. A higher perceived age was shown to be significantly associated with a lower bone mineral density (BMD)/trabecular bone score (TBS) in women aged 25-93 years [13].

Facial ageing is caused by intrinsic (genetic) and extrinsic (environmental) mechanisms. Multiple studies assess the impact of lifestyle like diet, sleeping, smoking, on facial aging [14], [15]. Further studies emphasize the importance of promoting psychosocial resources and mental health for the deceleration of facial aging as well as maintenance of good health [16].

Increasingly, facial aging studies correlate biology and perceived age, including specific facial features such as wrinkles, with profiling participants DNA by genotyping arrays and sequencing, and RNA. Several studies have utilized Genome Wide Association Studies (GWAS) to elucidate the effect of genetic factors on skin aging and specific aging features. One GWAS reported SNPs in three genes (DIAPH2, EDEM1, KCND2) associated with skin youthfulness as determined by a board-certified dermatologist from frontal facial photographs of participants [17].

Another study [18] analyzed whole transcriptomes of skin biopsies from the sun-protected inner arm area of 122 healthy women of European descent, ages 18-89 years with Fitzpatrick skin type I/II. The analysis was performed in the context of the perceived age as determined from digital photographs by the Canfield medical imaging software by four dermatologist raters blinded to the chronological age of the volunteers. An unbiased search for gene expression associated with intrinsic skin youthfulness, defined as the top 10% of individuals whose assessed skin aging features were most discrepant with their chronological ages, and identified several novel candidate genes including PHLdA1, HAS2-AS1. Additionally, this study demonstrated that immunologic gene sets are the most significantly altered in skin youthfulness, suggesting the immune system plays an important role in skin youthfulness.

An excellent recent discussion on genetic basis of skin youthfulness reviews how genetic variations, epigenetic changes, gene expression of RNAs and micro-RNAs, and mitochondrial depletion relate to skin aging [19].

In this paper, we report the results of functional analyses of genome-wide data on self-perceived facial aging by the participants of the UK Biobank (UKBB). It is well known that people often deceive themselves about their appearance [1]. Indeed, over 73% of people (of both genders) replied that other people say they “look younger than their age”, nearly 25% replied that others tell them they “look about the same age as they are”, and only 2% of participants replied that other people comment they look older. While the UKBB self-perceived facial aging data appears very skewed, we set to explore whether any insights into the biology of facial aging can be obtained from subjective answers to the question of how other people perceive your age. Surprisingly, we found the same biological pathways associated with self-reported facial aging as were reported by other studies where aging was carefully evaluated from images, measured, or assessed by specialists. Moreover, our results are in line with gene expression studies where the biological age of participants is known.

## Methods

### Data

In this paper we utilize results of GWAS conducted by the Neale lab http://www.nealelab.is/uk-biobank/ across thousands of traits, including facial aging, of 361K unrelated individuals in UK Biobank (UKBB). Over half a million participants from the UK Biobank have responded to the question “Do people say that you look” (1) “Younger than you are” (378,962), (2) “Older than you are” (10,306), (3) “About your age” (119,913) (available: http://biobank.ctsu.ox.ac.uk/crystal/field.cgi?id=1757).

We obtained genetic variations with suggestive associations by filtering downloaded genetic variations by p-value threshold=0.00001. Additionally, filtered out low confidence variants (*low_confidence_variant=TRUE*). For female participants, this results in 2352 SNPs that are used for further analysis. Taking the same p-value threshold for males, results in 8436 SNPs used for further analysis.

### Annotation

Genetic variations were annotated with dbSNP IDs, and also further annotated by genes (gene symbol) using *BiomaRt* package from R bioconductor. Additionally, each SNP was mapped to genes by using expression trait loci (eQTL) in skin (Skin - Sun Exposed, Lower leg, and Skin - Not Sun Exposed, Suprapubic) and cells-transformed fibroblasts from the *GTEx* portal (https://gtexportal.org).

#### Analysis

Enrichment analysis was performed *using XGR: eXploring Genomic Relations at the gene, SNP and genomic region level through enrichment, similarity, network and annotation analysis* (available: http://xgr.r-forge.r-project.org). Enrichment analysis was performed on all SNPs with suggestive associations. Additionally, enrichment analysis was performed on all SNPs that are in linkage disequilibrium (LD) with these SNPs (EUR population, with default R2 cutoff =0.8). Enrichment analysis on gene level was performed using an option with *Canonical/KEG/Biocarta pathways from MSigDB and Gene Ontology Biological Processes.*

## Results

### Genes known to be associated with facial aging

A recent comprehensive review on genetics of facial youthfulness [19] reports SNPs in genes that have been found to be associated with facial aging in several genome wide association studies (GWAS). Top category are the genes with roles in skin pigmentation (*MC1R, BNC2, RALY/ASIP, IRF4*). Specifically, melanocortin 1 receptor, *MC1R*, has been implicated in both global facial aging, as well as specific features of skin aging, including six-fold increased photoaging risk for carriers of loss-of-function variations and facial pigmented lesions [19-21].

For the exploratory analysis of the UKBB data on subjective facial aging, we interrogated a pool of SNPs with suggestive associations (p-values <=0.00001). This p-value threshold is higher than usually reported by GWAS for individual genetic variations. We nevertheless decided to explore whether genes, and pathways, previously reported in the context of aging are found in this dataset. Additionally, the gene space is extended by considering expression Quantitative Trait Locus (eQTL) that estimates the influence of a SNP’s presence on the expression of a gene in skin (Skin - Sun Exposed, Lower leg, and Skin - Not Sun Exposed, Suprapubic) and cells-transformed fibroblasts from the *GTEx* portal (https://gtexportal.org)

Indeed, several SNPs in *MC1R* (rs1805007; p-value=E-6) are associated with self-perceived facial aging with carriers of alleles responsible for lower pigmentation more likely reporting looking older than their actual age (S1 Table). SNPs in other pigmentation genes, previously found to be related to facial pigmentation spots [19], are also associated with self-reported aging: *IRF4* (16 SNPs with rs398000140 being the most significant, p-value=E-18), *RALY/ASIP* (rs6059655,rs1205312,rs6088372), *BNC2* (rs10810645), and *PIGU* (rs2424995). Additionally, over 30 eQTLs are in the *ASIP* gene in skin and fibroblasts (S1 Table).

A hallmark of aging skin is its loss of elasticity associated with damage to the extracellular matrix protein elastin, encoded by the ELN gene. Given the low turnover of elastin and the requirement for the long term durability of elastic fibres even relatively subtle mutations in the ELN gene may significantly impact the assembly and mechanical properties of human elastin [22].

One example is rs2071307 SNP, where carriers of the minor allele demonstrate improved performance from their elastin bearing tissues compared to the reference allele. Interestingly, the frequency of this allele in European populations is quite high (up to 40%) as stronger elastin provides various evolutionary advantages. The UKBB facial aging data for females reveals that three SNPs in the *ELN* gene (rs55863875, rs28763981, rs17855988) have suggestive associations with self-reported youthfulness. All three SNPs are in high LD (D’>=0.8) with the previously studied in [22] rs2071307 SNP but frequencies of the minor alleles for these SNPs are much lower. At least one of the SNPs, rs17855988 is missense and it is associated with cutis laxa, a rare disorder associated with deficient or absent elastin fibers in the extracellular matrix. While male self-reported facial aging data analysis does not have any variations in the ELN gene that are associated with facial aging, data for both sexes reveals 28 variations in the ELN gene associated with youthfulness (p-values<E-6).

Another gene known to influence skin aging via glycation is NAT2. It codes for the enzyme that participates in the detoxification of various drugs and toxins. In fact, polymorphisms in this gene segregate human populations into rapid, intermediate, and slow acetylator phenotypes. In the context of skin aging, SNPs in the NAT2 gene (rs1495741, rs1495714) are implicated in skin fluorescence that is a noninvasive measure of advanced Glycation End products (AGEs). Several SNPs in the NAT2 gene are significantly correlated with self-reported facial aging in the UKBB data.

### SNP enrichment analysis

We then set to investigate whether SNPs associated with self-perceived facial aging in females are enriched in specific functional categories and pathways. To this end, we ran the XGR analysis tool on all SNPs with suggestive associations (p-value<=0.00001; see Methods).

Nearly a third of over 70 enriched categories (FDR<=0.05) are related to skin traits, including skin diseases and inflammatory skin disorders. Top highly significant enriched categories are skin pigmentation (FDR=1.5E-23), suntan (FDR=2.7E-22) and skin sensitivity to sun (FDR=E-8). These results are in line with previously reported findings that pigmentation genes play crucial roles in skin aging [19]. Dyspigmentation is clinically associated with both intrinsic and extrinsic aging of skin and can be manifest as hyperpigmentation or hypopigmentation Indeed, as the number of melanocytes in our skin decreases and skin color in sun-protected areas lightens with age [23], aberrant pigmentation (hyper- and hypopigmentation; i.e., sun spots and freckles) occurs at chronically sun-exposed body parts [24].

The next highly significantly enriched categories are related to skin cancers and skin diseases, including rosacea (S2 Table). Several hair related categories are significantly over-represented, including hair morphology, hair color as well as alopecia. Indeed, hair loss is one of the hallmarks of perceived facial aging. Several categories related to appearance (abnormality of the mouth, lobe attachment, outer ear morphology trait) are also enriched.

Another large group of highly enriched categories are related to lung characteristics and diseases, including chronic obstructive pulmonary disease, susceptibility to pneumonia measurement, and even lung cancers. Lung disease (FDR=8.5E-38) and chronic obstructive pulmonary disease (FDR=2.4E-30) are also two top over-represented categories associated with increased facial aging in the analysis that includes SNPs in high LD (EUR population, R2=0.8; S3 Table)). Additionally, other lung related categories such as forced expiratory volume, emphysema, respiratory system neoplasm are associated with accelerated facial aging. Several studies have demonstrated a common susceptibility of lung and skin aging [25]. Interestingly, several categories related to smoking (FDR=2E-8) are also over-represented in SNPs associated with accelerated self-reported facial aging. This is in line with numerous studies that have indicated correlations between skin wrinkling/aging and smoking.

An interesting observation is that variations found in GWAS to be related to mosquito bites (attractiveness to mosquitoes and itch size) are significantly over-represented in the data. However, the direction of the association between these two traits is not clear, partly due to ambiguous outcomes for some variations (e.g. rs521977 is reported to being related to both increase in the itch size and and at the same time to increase in unattractiveness to mosquitoes). Still, it appears that some variations and genes related to mosquito bites are implicated in facial aging, and this may be of interest to be investigated further.

### Enrichment analysis on gene level

We then ran XGR enrichment analysis on annotated (284) genes using the option *canonical/KEGG/REACTOME/Biocarta*. Separately, we ran enrichment analysis on genes mapped by using expression trait loci (eQTL) in skin or fibroblasts (162 genes; see Methods). These will be referred to as eQTL skin-expressed genes.

The majority of enriched pathways are related to the immune system (S4 Table). Immune related pathways are also over-represented in eQTL skin-expressed genes (S5 Table). This is in line with two gene expression analyses [18], [26]. Immune response and cell adhesion were identified among mechanisms associated with gene expression changes in aged skin versus young skin [26]. Whole transcriptome profiling revealed the immunologic gene set as the most significantly altered in people with skin youthfulness versus others, identifying the immune system as a potential contributor to youthful appearing skin [18]. Dysregulation of immune system genes with aging has been reported as one of the six gene expression hallmarks of cellular ageing in a survey of age-linked changes in basal gene expression across eukaryotes from yeast to human [27].

Other significantly overrepresented gene-sets related to skin aging include semaphorin interactions, structural ECM glycoproteins, cell adhesion molecules, transport of glucose and other sugars, bile salts and organic acids, metal ions and amine compounds, and (marginally significant) biological oxidations (S4 Table). Nearly all these categories have been implicated in skin aging, or wound healing (semaphorins) [26]-[29].

Interrogating biological pathways (GOBP) yields overrepresentation of age-related cellular response to epidermal growth factor stimulus, and fibroblast growth factor, as well as responses to heat, nutrients, oxidative stress, and inflammation (S6 Table), in addition to housekeeping cellular processes, such as protein folding, phosphorylation, and mRNA splicing.

Functional analysis of biological pathways in the list of eQTL skin-expressed genes reveals enrichment of several additional immune-related pathways, and pathways related to metabolism of lipids, lipoproteins and fatty acids (S7 Table) in addition to 14 over-represented pathways that are common for both gene lists. Among other over-represented biological processes in eQTL skin expressed genes, it is worth mentioning apical junction complex, and related adherens junction organization, cell-cell adhesion, transport of sialic acid and sodium, cellular response to glucose stimulus and inflammation, metabolic processes for urate, carbohydrate, and cellular protein, lipid biosynthesis, and several immune related processes.

### Comparison between self-perceived female and male facial aging

As females and males perceive aging differently, it is perhaps not surprising that the genetic correlates with self-reported facial aging do not overlap much on the level of individual SNPs, or genes. There are over 3-times more SNPs associated with the male responses than those with female responses (for the same p-value thresholds). Can it reflect a slightly more objective male responses compared to self-perceived youthfulness by females? Only a small fraction (12%) of SNPs (292), and 24 genes, overlap for both genders. These common SNPs are significantly enriched in skin and hair pigmentations, and skin diseases categories, while the only over-represented category in the common genes list is related to extracellular matrix (FDR=0.009; *EFEMP1, LOXL1, PLXNA1*).

Similar to females, functional analysis of males’ facial aging reveals overrepresentation of skin pigmentation traits (FDR=E-10) as well as hair-related traits, including hair loss. Figure 1 shows comparison of enrichment results for female(in red) and make (in blue) using side-by-side barplots displayed by -log10(FDR). In fact, hair loss (alopecia) is among the top three over-represented categories for males (FDR=E-59; S8 Table). Interestingly, over 10 categories related to measurements of lipids and fatty acids (high density lipoprotein cholesterol, cis/trans-18:2 fatty acid, trans fatty acid, omega-6 polyunsaturate, linolenic, linoleic, docosapentaenoic acids, lipoproteins) are overrepresented in males facial aging data, and correlated with self-perceived facial youthfulness.

**Figure 1:**
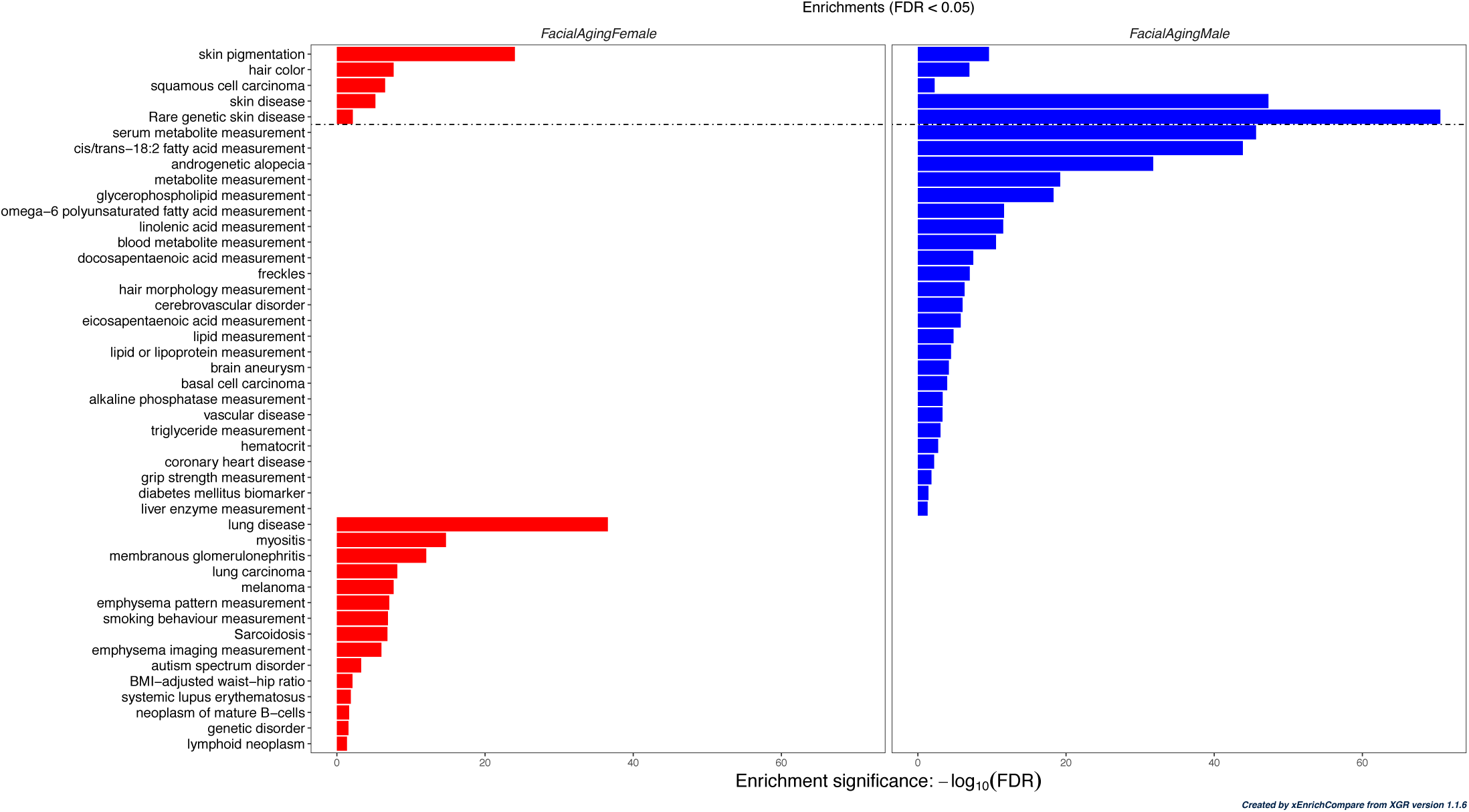
Visualization of enrichment analysis results for SNPs associated (FDR<E-5) with female (red) and male (blue) self-perceived facial aging. Enrichment is run on previously identified SNP-based traits/categories (Experimental Factor Ontology), taking into consideration SNPs that are in LD (EUR, r2=0.8). Traits/categories are displayed by -log10(FDR) with FDR<0.05. Enrichment analysis is performed using XGR R library, and visualized using xEnrichCompare() function.

Functional analysis on gene level confirms enrichment of lipid-related categories, including steroid hormone biosynthesis, metabolism of lipids and lipoproteins, Fatty acid, triacylglycerol, and ketone body metabolism (S9 Table, Figure S1). At least one study [30] revealed reduced expression of genes related to lipid biosynthesis in both chronologic and photoaged skin by comparing transcriptomic data from sun exposed and sun protected skin in young and old individuals. Additionally, functional analysis of genes related to self-perceived male aging reveals several signaling pathways (e.g. TGF-beta signaling pathway) and cell-cycle pathways (e.g. Cell Cycle: G1/S Check Point) that are not observed in the self-perceived female aging data.

### Next steps: identifying novel biomarker candidates from self-reported facial aging data

Among common pathways significantly overrepresented in the data for both genders is the gene-set for semaphorin interactions. Semaphorins are known to protect tissue barriers, and regulate epithelial and endothelial junctions [31]. Specifically, PLXNA1 (Plexin-A1), the only overlapping gene in the data for both genders in the semaphorins interactions category, belongs to the family of plexins that are the main cell surface receptors for semaphorin signal transduction. There are multiple genetic variations in PLXNA1 gene associated with facial aging responses by both females and males. Moreover, dozens of eQTLs for Plexin-A1 in female data, and over 150 eQTLs in male data, are significantly associated with aging in skin expressed genes in skin or fibroblasts. While semaphorins are known to play roles in cell migration and motility, wound healing, modulation of skin inflammation and sensitivity, specific role of Plexin-A1 in skin aging has not been reported, and it warrants further investigation.

Structural glycoproteins of the extracellular matrix (ECM) play an important role in skin aging. Wrinkling, or loss in skin elasticity, results from matrix metalloproteinases (MMPs), key components of the ECM, degrading the main constituents of the ECM such as types I and III collagen, proteoglycans and glycosaminoglycans (GAGs). Also, the amount of elastin decreases with age. In this analysis, structural ECM glycoproteins were found to be significantly enriched in both female and male data on facial aging. This aligns with two gene expression studies that have identified structural ECM glycoproteins as significantly associated with skin aging [18], [32].

While the ECM genes implicated in the distinct datasets vary, the only overlapping gene present in the UKBB facial aging data for both genders, as well as two gene expression studies - is EFEMP1 (Fibulin-3; S10 Table). EFEMP1 is significantly upregulated with aging in both gene expression studies referenced above. In alignment with this upregulation, our analysis found over 40 SNPs in the EFEMP1 gene that are associated with self-reported accelerated facial aging (“looking older”). EFEMP1 has also been reported to be overexpressed in senescent human fibroblasts that accumulate in skin with age [33], [34].

## Discussion

With skewed answers, indicating that most respondents self-reported that they are perceived by others to be younger than they actually are, one would initially assume the data to be disorganized noise, uncorrelated with any underlying biology and genetic factors. However, the surprising results of this analysis reveal that, in fact, the self-perceived data on facial aging provides several interesting biological insights. By interrogating correlations between self-reported facial aging and hundreds of publicly available traits and pathways, we identified several age-related biological pathways and categories that are over-represented in the data. Furthermore, the enriched categories align with previous studies that used standardized measures of facial aging and found similar or identical genetic influences on facial aging.

Specifically, consistent with earlier observations pigmentation genes and pathways are correlated with facial aging, quantified from imaging, perceived by others, or self-reported. Skin cancer is one of the top over-represented categories associated with facial aging. Interestingly, a recent independent study on another cohort (Leiden Longevity) found that mechanisms influenced by genetic loci increasing susceptibility to basal cell carcinoma also drive skin ageing suggesting shared biology and shared targets for interventions [35].

Additionally, numerous immune-related gene-sets and pathways significantly associated with facial aging data as has already been found in gene expression studies that systematically compared transcriptomes from young and old skin samples.

One of the interesting findings from this data is further validation that structural ECM glycoproteins are significantly correlated with facial aging. This aligns with results from two independent gene expression studies [18, 19, 32]. All three studies identified fibulin-3 (*EFEMP1*) as one of the key proteins related to skin ageing. As *EFEMP1* is overexpressed in senescent fibroblasts that increase with age, it is a good candidate for further exploration as a marker of skin senescence and aging. Another plausible candidate that emerges from this analysis is plexin-A1 (*PLXNA1*) that is known to play a role in axon guidance, invasive growth and cell migration but has not yet been implicated in facial aging.

It is worth noting that less than 5% of the UKBB participants reported “looking older than actual age.” These numbers are very much in line with results from another study of over 4,000 participants from Australia, Canada, and the UK, where nearly all participants (95.6%) perceived themselves as “looking your current age or younger” [36].

While it is a standard approach that GWAS report genetic variations results with significance of p-values<E-8, using such strict threshold will result in missing many loci of potential interest. As self-perceived facial aging is a very complex trait, influenced by genetics, environment (from skin care to sun exposure and lifestyle) as well as psychology, and social interactions, in this study we considered genetic variations with suggestive associations(p-values<E-5). With the understanding that some identified results are false-positives, it is very likely that additional genetic signals exist beyond the ones captured and explored in this study.

Overall, we demonstrate that re-analysis of large-scale genotyping data related to facial aging sheds light on pathways of skin aging, and even reveals age-related biomarkers. Moreover, our study re-confirms a large genetic component in facial aging. We believe that collection of large-scale data on facial aging using self-reported surveys, imaging data, and genetics will soon pave the way to identifying skin biomarkers of facial aging leading to truly personalized approaches in dermatology.

## Supporting information

**S1 Table. Annotated genetic variations associated with self-perceived female facial aging.**

Subset from the UKBB data for female facial aging with p-values<=0.00001. Data downloaded from http://www.nealelab.is/uk-biobank. Annotated with dbSNP IDs, genes (using BioMart package from R Bioconductor), and eQTLs in skin (Skin - Sun Exposed, Lower leg, Skin - Not Sun Exposed, Suprapubic) and cells-transformed fibroblasts from the GTex portal (https://gtexportal.org)

**S2 Table Enrichment analysis on SNP level for female facial aging data**

XGR: SNPs based enrichment analysis. Experimental Factor Ontology. Female facial aging data. Not including LD SNPs.

**S3 Table Enrichment analysis on SNP level (including SNPs in high LD) for female facial aging data**

XGR: SNPs based enrichment analysis. Experimental Factor Ontology. Choose population (from 1000 Genomes Project) for which to include SNPs in Linkage Disequilibrium (LD) with GWAS lead SNPs. EUR population (LD, R2 cutoff=0.8). Female facial aging data.

**S4 Table Enrichment analysis on gene level for annotated genes using pathways ontology for female facial aging data**

XGR gene-based enrichment analysis. Ontology:

canonical/KEGG/REACTOME/Biocarta pathways. Female facial aging data.

**S5 Table Enrichment analysis on gene level for eQTL genes using pathways ontology for female facial aging data**

XGR gene-based enrichment analysis. Skin expressed genes (genes mapped by using expression trait loci (eQTL) in skin or fibroblasts). Ontology:

canonical/KEGG/REACTOME/Biocarta pathways. Female facial aging data.

**S6 Table Enrichment analysis on gene level for annotated genes using biological processes ontology for female facial aging data**

XGR gene-based enrichment analysis. Ontology: Biological Processes. Female facial aging data

**S7 Table Enrichment analysis on gene level for eQTL genes using biological processes ontology for female facial aging data**

XGR gene-based enrichment analysis. Skin expressed genes (genes mapped by using expression trait loci (eQTL) in skin or fibroblasts). Ontology: Biological Processes.

Female facial aging data.

**S8 Table Enrichment analysis on SNP level for male facial aging data**

XGR: SNPs based enrichment analysis. Experimental Factor Ontology. Male facial aging data.

**S9 Table Enrichment analysis on SNP level (including SNPs in high LD) for female facial aging data**

XGR gene-based enrichment analysis. Ontology:

canonical/KEGG/REACTOME/Biocarta pathways. Male facial aging data.

**S10 Table.** Extracellular matrix genes found associated with aging in three independent studies. Extracellular Matrix (ECM) genes and their overlap in three studies: UKBB facial aging (genotyping data), whole transcriptome sequencing of skin biopsies from sun-protected inner arm of older women with skin youthfulness compared to older women without skin youthfulness [18], and gene expression levels changes with age in abdominal skin from 856 female twins [32]. Gene overlap between data sets is shown in **bold** font.

| Xu et al [18] | Glass et al [32] | UKBB facial aging |
| --- | --- | --- |
| *CILP, CRISPLD2, CTHRC1, DPT, **EFEMP1**, FBLN2, FBLN5, FBN1, LAMC1, LTBP2, **MFAP4**, MGP, MMRN1, POSTN,* | *COLQ, **EFEMP1**, LAMB3, LAMB4, NTN5, PAPLN* | ***EFEMP1**, ELN, MATN2, **MFAP4**, TNXB, VWA7* |
| *SLIT3, SVEP1, THBS2,*<br>*VWA5A* |  |  |

